# Use and abuse of potential rates in soil microbiology

**DOI:** 10.1101/2020.05.18.101253

**Authors:** Christina Hazard, James I. Prosser, Graeme W. Nicol

## Abstract

Potential rate assays are used in soil microbial ecology to determine the rates of a functional process in environmental samples under a defined set of conditions. While they can be used appropriately to provide mechanistic insights, potential rates are also often used to estimate the abundance of specific taxonomic groups and their *in situ* activity. These estimates incorrectly assume that all contributing organisms in a community are active at a maximum rate under one set of ‘optimal’ incubation conditions and that potential rates reflect activity in the soil. While investigators now recognise that populations within communities are physiologically diverse, they often ignore the consequent suboptimal activity, or even inactivity, of the majority of community members performing that function. In this short perspective article, we discuss when potential assays can be informative and highlight the underlying conceptual problems under circumstances where potential assays are misused, using potential nitrification rate (PNR) as an example. PNR was originally developed to estimate the size of active ammonia oxidising communities in environmental samples. It is routinely determined in short-term shaken slurry incubations by measuring assumed maximum rates of nitrate or nitrite production under optimal, non-substrate-limiting conditions. As with other functional processes, it is now recognised that a broad diversity of organisms contribute to aerobic ammonia oxidation in terrestrial and other habitats, and this diversity represents a substantial range of physiologies, including variation in substrate affinity, ammonia tolerance, cell specific activity and substrate preference. Despite this, PNR, and other potential rate assays, are often inappropriately used in an attempt to determine an ecologically relevant measurement of activity in soil. As with any potential assay, PNR has inherent biases towards particular functional groups and its use in investigating the ecology of ammonia oxidisers in natural systems should be carefully considered.

## Main text

Measurement of the potential rate of microbially mediated soil processes has a long tradition. Commonly used examples are measurement of potential nitrification and denitrification rates and substrate-induced respiration. All determine a process rate, usually after addition of a relevant substrate, under what are sometimes considered to be optimal conditions that enable most organisms capable of performing the process to operate with maximum activity and/or specific growth rate. Potential rate obviously differs from ‘actual’ process rate, which will typically operate under non-optimal environmental conditions. Potential process rates have value, and have traditionally been used, in the absence of alternatives, to estimate the biomass (or, less justifiably, abundance) of a functional microbial group. However, the assumptions underlying measurement of potential rate are often not considered, obeyed or tested and have been increasingly challenged as our understanding of the composition and diversity of microbial communities in natural environments has increased. The value, limitations and relevance of potential rate measurements in microbial ecology are illustrated below for soil nitrification. However, the factors defining the usefulness and limitations of potential assays will also apply to other processes and functional groups, some of which encompass far greater taxonomic or functional diversity. For example, potential denitrification assays are typically performed using one set of incubation condition only and attempt to target organisms that represent a far greater level of taxonomic and physiological diversity than nitrifiers. In such circumstances, predictive power will likely be even further reduced.

### Potential nitrification rate

Nitrification is the sequential oxidation of ammonia (NH_3_) to nitrite (NO_2_^−^) and nitrate (NO_3_^−^) and is an essential component of reactive nitrogen transformation in soil. While nitrite can accumulate in soil (Giguere et al., 2017; Ma et al., 2015; Taylor et al., 2019; Venterea and Rolston, 2000), particularly after N deposition (Venterea et al., 2020), ammonia oxidation is often assumed to be the rate-limiting step. As a consequence, the activity, diversity and community structure of ammonia oxidisers (AO) receives greatest attention with respect to understanding environmental factors that influence microbially-mediated nitrification. Potential nitrification rate (PNR) assays (also referred to as potential nitrification activity (PNA)) have been used for many decades as a rapid method for estimating net ammonia oxidation (≈nitrification) rates in soils.

While a range of PNR assays have been used, the most common approach is the shaken soil-slurry method (Belser and Mays, 1980; Hart et al., 1994). This typically involves incubation of a suspension of sieved soil in a solution containing NH_4_^+^, at non-limiting concentration, in a shaken flask, ensuring aeration and mixing, at an incubation temperature that is considered to be optimal. PNR is calculated as the gradient of the linear increase in nitrate concentration in samples taken regularly during a relatively short incubation period (often ≤24 h) (Hart et al., 1994). A potassium phosphate buffer, typically between pH 7.0 and 7.5, can be included in the assay solution, with the intention of optimizing pH for AO. Chlorate, an inhibitor of nitrite oxidation, is also commonly included and NO_2_^-^ rather than NO_3_^-^ is measured in each slurry sample (Belser and Mays, 1980). Growth is assumed to be negligible; hence the assumption of a linear, rather an exponential increase in nitrite or nitrate concentration. PNR therefore measures the activity of the community and not specific growth rate. Linear kinetics are often not tested and it is not unusual for calculations to be based on nitrate concentrations measured only at the beginning and end of incubation or even using just the final value. Any growth, or other divergence from linear kinetics, invalidates calculation of a ‘constant’ PNR.

Employment of ‘ideal’ or ‘optimal’ conditions attempts to ensure that all AO within the sample are active, which is further encouraged by addition of substrate (NH_4_^+^). The assumption of universal activity was implicit in the traditional use of PNR to estimate AO abundance in soil. This assumption was always seen as a major limitation of this approach (Woldendrop and Laanbroek, 1989; Hart et al., 1994), particularly given the low growth rates of ammonia oxidisers. (The only alternative was use of the most probable number method, which also relies on incubation of dilutions of soil suspensions under ‘ideal’ conditions and similar assumptions.)

### Physiological diversity of ammonia oxidisers challenges underlying assumptions of PNR

For more than a century, ammonia oxidation was thought to be performed mainly by ammonia oxidising bacteria (AOB) (Prosser and Nicol, 2012). The past 25 years have, however, demonstrated considerable phylogenetic diversity within AOB and also the capacity for ammonia oxidation by ammonia oxidising archaea (AOA) (Könneke et al., 2005) and complete ammonia oxidizers (comammox) of the genus *Nitrospira* (Daims et al., 2015; van Kessel et al., 2015). Evidence from both cultivation-based and ecological studies also provides evidence for substantial physiological diversity in AO. Physiological diversity of cultivated AOB is well established and was used for traditional classification of these organisms (Prosser 1989; Koops et al., 2003). Since the first isolation of an ammonia oxidising archaeon (AOA), *Nitrosopumilus maritimus* (Könneke et al., 2015), several enrichments and isolates of soil AOA have also been obtained. These include, for example, the neutrophile *Nitrososphaera viennensis* EN76 (Tourna *et al*. 2011) and the obligate acidophile *Candidatus* Nitrosotalea devanaterra ND1 (Lehtovirta-Morley *et al*., 2011). In addition, comammox *Nitrospira* are abundant and contribute to ammonia oxidation in soil (Pjevac et al., 2017; Wang et al., 2019a), and a gammaproteobacterial AOB (*Candidatus* Nitrosoglobus terrae) was isolated recently (Hayatsu *et al*., 2017). This taxonomic diversity is reflected in ecophysiological diversity and the maximum specific growth rate, cell specific activity, inhibitory ammonia and nitrite concentrations and affinity for ammonia of AO all vary by over an order of magnitude among different soil isolates, both within and between the major ammonia oxidiser groups (summarised in Lehtovirta-Morley et al., 2016). In addition, many AO are inhibited by NO_2_^-^, which accumulates when chlorate is used to inhibit nitrite oxidisers, thereby reducing PNR depending on community composition. Thus, even if the incubation conditions (temperature, pH, oxygen concentration) are appropriate for all AO, the PNR will represent only an aggregate of a wide range of cell activities and will vary with community composition and not just abundance.

Physiological diversity does, however, also mean that incubation conditions required for activity of all AO will not be identical. Cultivation-based studies have demonstrated significant differences in effects of temperature (Jiang and Bakken, 1999; Tourna et al., 2011; Lehtovirta-Morley et al., 2014), pH (Allison and Prosser, 1991; Lehtovirta-Morley et al., 2011) and salinity (Mosier et al., 2012; Qin et al., 2014; Bello et al., 2019). Similar evidence of physiological diversity is provided by ecological studies (Nicol et al., 2008; Ollivier et al., 2012; Gubry-Rangin et al., 2017; Nacke et al., 2017; Taylor et al., 2017; Lu et al. 2018) and the ubiquity of AO in the full range of soils, in itself, is evidence that a single set of incubation conditions will be insufficient. For example, ammonia supply is an important factor influencing niche differentiation between AOA and AOB populations in soil. AOB often dominate growth at higher concentrations of added inorganic ammonium (e.g. Jia and Conrad, 2009; Xia et al., 2011; Hink et al., 2017), while AOA appear to prefer ammonium that is derived from a mineralised organic N source, i.e. that is part of the native organic component of the soil or added organic N (Levičnik-Höfferle et al., 2012; Hink et al., 2018). In low pH soils, AOA make a greater relative contribution to ammonia oxidation under specific incubation conditions (Gubry-Rangin et al., 2010; Zhang et al., 2012) and, as such, a decreasing proportion of the ammonia oxidising community may preferentially oxidise added inorganic N with decreasing soil pH.

It is therefore impossible to identify incubation conditions that will be ‘ideal’ for all organisms and the majority of AO will have sub-optimal or no activity and will not contribute to measured PNR. To this must be added the assumption that ammonia available and oxidised in PNR assays will also be available for the same populations in soil.

### *Potential nitrification rate does not reflect ammonia oxidiser activity* in situ

The consequences of these limitations can be seen when correlations are determined between published PNR for soils and sediments and AOA and AOB abundances (using *amoA* gene abundance as a proxy for cell abundance). Results are highly variable. Of 107 studies published from 2007 to 2020 (Supplementary Table 1), only 36% and 75% reported a significant positive correlation between PNR and AOA or AOB abundances, respectively (Figure 1). Ten percent and 46% report a positive correlation with only AOA or only AOB, respectively, 11% correlated with neither and 4% and 5% reported a negative correlation with AOA and AOB, respectively. No correlation has yet been observed between PNR and comammox bacteria (Wang *et al*. 2019b; Luchibia *et al*. 2020; Zhang *et al*., 2020). While these studies suggest that some populations of AOA are using added ammonium, the majority of studies indicate that only ammonia oxidising bacteria (AOB) contribute to PNR, often leading to the conclusion that AOA are not contributing to ammonia oxidation in a soil *in situ*. If we consider the growth characteristics of AO in both culture and soil, it is evident that correlating results from a single defined assay with the abundance of communities containing substantial physiological diversity is unlikely to be meaningful and likely to be misleading.

**Figure 1.** Meta-analysis of soil and sediment studies finding significant positive correlations of AOA and/or AOB abundances with PNR. All studies used a shaken slurry method and measured NO_3_^-^ or NO_2_^-^ with the addition of chlorate to inhibit NO_2_^-^ oxidation. A total of 107 studies were included (Supplementary Table S1).

The observed dominance of AOB over AOA in contributing to PNR activity in many soils may reflect differences in their ability to utilise added inorganic nitrogen. To investigate this, soil was sampled across a pH 4.5 to 7.5 gradient that has been fully characterised with respect to AOA and AOB community structure and abundance (Aigle et al., 2019; Gubry-Rangin et al. 2011), where *amoA* transcript and gene abundance of AOA and AOB decrease and increase directly with soil pH, respectively (Prosser and Nicol, 2012). PNR assays were established with both standard 1 mM phosphate buffered and unbuffered 1.5 mM NH_4_^+^ solution, and parallel incubations were performed in the presence of alkynes to differentiate contributions of all AO, AOA and AOB. Acetylene (C_2_H_2_) has been extensively used as an inhibitor of all ammonia oxidisers that possess ammonia monooxygenase (AMO) and 1-octyne (C_8_H_2_) inhibits growth of AOB, but not AOA, in both culture and soil (Taylor et al., 2013). Nitrate accumulation during incubation for 24 h was used to calculate PNR, which increased directly with soil pH across the gradient (Figure 2). However, as this activity was completely inhibited in the presence of both C_8_H_2_, and C_2_H_2_, it was attributed solely to AOB. This provides strong evidence that AOA do not contribute to PNR in some soils, PNR severely underestimates their abundance and this prevents any meaningful interpretation of their abundance or actual activity in soil at the time of sampling.

**Figure 2.** Demonstration of only AOB contributing to PNR. Soils were sampled across a pH gradient and PNR determined using the shaken slurry method. Flasks were established in triplicate using 144 ml serum vial bottles with 6 g soil (dry weight equivalent) and 50 ml of 1.5 mM NH_4_Cl, with or without 1 mM potassium phosphate (pH 7.2) buffer. (a) Samples were taken after incubation for 0, 3, 6, 9, 12 and 24 h to determine nitrate concentrations (nitrite was not detected). Means (±SE) with different letters within a buffer treatment are significantly different based on Tukey’s *post hoc* test following one-way ANOVA (*p*<1.1 × 10^−7^). Means annotated with a line are significantly different between buffer treatments based on two sample *t*-tests (*p*>0.01<0.05). (b) Example of complete inhibition of nitrification activity in flasks incubated in the presence of 0.01% C_2_H_2_ or 0.03% C_8_H_2_ with pH 7.5 soil (unbuffered). Nitrification activity was not detected in any soil in the presence of either alkyne.

As indicated above, the source of ammonium is also an important factor in defining the ecological niches of AO populations within a community. Using concentrations typical of fertilisation application rates, we compared nitrification activity in both pH 4.5 and 7.5 soil microcosms after the addition of ammonium nitrate or urea (which is fully mineralised to ammonium within 0.5 days). After near stoichiometric conversion of ammonium to nitrate over 12 days, inorganic and urea-derived ammonium were oxidised at comparable rates for both 100 and 200 µg N g^-1^ dry soil applications in the pH 7.5 soil (Figure 3). The greatest change in pH in soil microcosms was associated with those with the highest nitrification rates, indicating that pH change was not an inhibiting factor (Supplementary Figure 1). Nevertheless, these results highlight an additional concern, as changing conditions during incubation will influence activity and kinetics will no longer be linear. In contrast, urea-derived ammonium was oxidised at a faster rate in the pH 4.5 soil, and ammonia oxidation was inhibited by application of 200 µg inorganic N g^-1^ dry soil (Figure 3). While these data do not differentiate the contribution of different AO groups, they reveal that sources of N other than inorganic NH_4_^+^, as used in PNR, are preferred by some AO populations, again preventing meaningful interpretation of PNR data.

**Figure 3.** Nitrate (NO_3_^-^) production in soil microcosms amended with varying concentrations of ammonium sulphate or urea (0 – 200 ug N g^-1^ dry soil) in pH 7.5 and 4.5 soil ((a) and (b), respectively). Soil microcosms were prepared (in triplicate for each treatment and timepoint) with sterile 144 ml serum vial glass bottles using 10 g soil (dry weight equivalent) which was sampled at 0-10 cm depth and sieved (2 mm mesh size). Microcosms were incubated at 30°C for 12 days with 30% soil water content (w/w). Microcosms were aerated every 3 days and destructively sampled before extracting NO_3_^-^ in a 1 M KCl solution. Nitrate concentration was determined colorimetrically following Hink et al. (2018) immediately after sampling (NO_2_^-^ was not detected). Error bars represent standard error of the mean. The greatest change in pH in soil microcosms was associated with those with the highest nitrification rates, indicating that pH change was not an inhibiting factor (Supplementary Figure 1). Nevertheless, these results highlight an additional concern, as changing conditions during incubation will influence activity and kinetics will no longer be linear.

These two experiments, performed using only one experimental agricultural soil system, are clearly not meant to be considered representative of all other managed or natural soil systems. They do, however, demonstrate that PNR does not effectively characterise the contribution of all AO populations to nitrification, but selects for those that are most competitive under one set of incubation conditions. Despite the major limitations discussed above, potential process rates are still frequently used inappropriately to address ecological questions. Attempts to understand the relative contribution of AOA and AOB to nitrification often involves correlation of their abundances with PNR, often after a treatment or perturbation (e.g. fertiliser application or temperature change) (Supplementary Table 1). This approach is also used to study niche specialisation, which implicitly assumes that changes in relative abundance of different phylotypes will result from differences in activity under different conditions. Incubation conditions used to measure PNR will favour different members of the AOA and AOB communities and will differentially influence activities of AOA and AOB. It will therefore not be possible to determine whether relationships between abundance and PNR are due to differences in preferences of the different communities for the specific incubation conditions used or due to links between abundance and actual activity. A more fundamental issue is the assumption, implicit in niche specialisation, that changes in community composition result from differences in actual activity, determined by different *in situ*, environmental conditions and do not depend on potential activity under a single set of laboratory conditions.

### When can potential rate measurements be useful?

Potential functional assays aim to employ incubation conditions that enable maximum or optimal activity for all organisms performing a particular microbial function in an environmental sample. They therefore provide a means of estimating the biomass of functional groups and standard conditions have been developed for such assays. This is exemplified by traditional use of potential nitrification rate, potential denitrification rate and substrate induced respiration as measures of biomass of nitrifiers, denitrifiers and ‘total’ microbial biomass in soil. This article has described the major disadvantages of this approach and such assays have very limited value as measures of biomass. Alternative methods are available and have largely been replaced for many functional groups by estimates of functional gene abundance. While these have their own limitations, they do not rely on or describe activity.

These traditional potential assays must, however, be distinguished from the use of laboratory incubations, in liquid culture, soil slurries or soil microcosms to study soil microbiology. Many such studies measure activity following substrate amendment and under controlled incubation conditions. Indeed, such studies are often used to investigate the effects of substrate type and concentration and environmental conditions on functional groups, and subgroups, to distinguish their activities and to investigate the influence of soil conditions. While such studies effectively measure ‘potential’ activity, this term is rarely used and is never used to imply that all organisms will be active or that the activity of each is optimal and identical. Laboratory incubations are ecologically meaningful if the limitations of the approach are critically evaluated with respect to their ability to assess activities in the environment (Taylor et al., 2019). For example, with respect to nitrification, they have demonstrated differences in the optimum temperature of AOA and AOB activity (Taylor *et al*., 2017), ammonia source preference (Levičnik-Höfferle *et al*., 2012) and nitrification-associated N_2_O yields (Hink *et al*., 2017). For other functional groups they have, for example, revealed the presence of both high- and low-affinity methane oxidisers in soil (Bender and Conrad, 1992) and greater influence on denitrification by edaphic factors than N-reducing community composition (Attard *et al*., 2011). In these instances, it is the relative changes in activity that provide valuable insights into the ecophysiology of different functional groups and not inference of a direct measurement of activity in the natural environment.

The use of *ex situ* potential activity assays for inferring *in situ* activity is not justified and better measures are available. For example, soil nitrification activity can be measured as gross nitrification rates using ^15^N pool dilution methods (Hart et al., 1996). These techniques may be more time-consuming and require specialist equipment, unlike PNR assays, but they provide environmentally relevant results. Where PNR has been used to evaluate the effect of an environmental factor, observations of relative differences in PNR will reflect effects on a particular set of nitrifying populations, and the data must be recognised as only reflecting the changing activities of these populations and not an overall community response. This is, of course, the same situation for any functional group utilising a specific substrate, but curiously this is often ignored. PNR does not characterise the contribution of all AO to nitrification in soil and knowledge of the diversity of AO physiology and factors controlling their growth clearly challenges a long-established assumption that a single incubation assay measures the activity of all AO. Critically, potential rates can have little relevance to actual rates *in situ* and therefore have limited meaning in many nitrification studies which should discourage the use of PNR in many circumstances.

Our expanding knowledge of the vast diversity and composition of microbial communities in natural environments emphasises that communities contributing to a functional process comprise a diverse array of ecophysiologies with substantial niche differentiation between populations. This challenges the underlying assumptions in using traditional potential rate measurements to estimate microbial biomass or *in situ* activity. However, controlled and manipulated laboratory studies that consider the underlying assumptions implicit in their use have been, and will continue to be, an important tool for mechanistic studies of soil microbiology.

## Acknowledgements

The authors would like to thank the reviewers for their insightful and constructive comments on the manuscript. This work was funded by the AXA Research Fund.

## Figures

**Supplementary Figure 1**. Soil pH in soil microcosms amended with varying concentrations of ammonium sulphate or urea (0 – 200 ug N g-1 dry soil) in pH 7.5 and 4.5 soil ((a) and b), respectively). The standard errors of the mean pH of triplicate microcosms were ≤0.06 and are not visible behind markers.

**Supplementary Table 1**. Analysis of 107 studies correlating PNR with the abundance of AOA, AOB or comammox bacteria as (inferred from qPCR analysis of *amoA* gene abundance). Published studies were found with a SCOPUS search (TITLE-ABS-KEY ammonia AND oxidi* AND archaea AND bacteria AND potential AND nitrification) combined with manual refinement for only soil and sediment studies that used a shaken soil-slurry method. Publications are grouped by soil or sediment and NO_2_^-^ or NO_3_^-^ measured as the end product. The majority of correlations were positive (X) although nine negative correlations (-X) were found. There were no correlations with comammox bacteria.

